# Species composition and altitudinal distribution of bumble bees (Hymenoptera: Apidae: *Bombus*) in the East Himalaya, Arunachal Pradesh, India

**DOI:** 10.1101/442475

**Authors:** Martin Streinzer, Jharna Chakravorty, Johann Neumayer, Karsing Megu, Jaya Narah, Thomas Schmitt, Himender Bharti, Johannes Spaethe, Axel Brockmann

**Affiliations:** Department of Neurobiology, Faculty of Life Sciences, University of Vienna, Vienna, Austria; Department of Zoology, Rajiv Gandhi University, Itanagar, Arunachal Pradesh, India; Obergrubstrasse 18, 5161 Elixhausen, Austria; National Centre for Biological Sciences, Tata Institute of Fundamental Research, Bengaluru, India; Department of Animal Ecology and Tropical Biology (Zoology III), Biocenter, University of Würzburg, Würzburg, Germany; Department of Zoology and Environmental Sciences, Punjabi University, Patiala, Punjab, India; Department of Behavioral Physiology and Sociobiology (Zoology II), Biocenter, University of Wurzburg, Wurzburg, Germany

**Keywords:** Conservation, Apidae, pollination, alpine habitats, global change, insect collection

## Abstract

The East Himalaya is one of the world’s most biodiverse ecosystems. Yet, very little is known about the abundance and distribution of many plant and animal taxa in this region. Bumble bees are a group of cold-adapted and high altitude insects that fulfill an important ecological and economical function as pollinators of wild and agricultural flowering plants and crops. The Himalayan mountain range provides ample suitable habitats for bumble bees. Himalayan bumble bees have been studied systematically for a few decades now, with the main focus on the western region, while the eastern part of the mountain range received little attention and only a few species are genuinely reported. During a three-year survey, we collected more than 700 bumble bee specimens of 21 species in Arunachal Pradesh, the largest of the north-eastern states of India. We collected a range of species that were previously known from a very limited number of collected specimens, which highlights the unique character of the East Himalayan ecosystem. Our results are an important first step towards a future assessment of species distribution, threat and conservation. We observed clear altitudinal patterns of species diversity, which open important questions about the functional adaptations that allow bumble bees to thrive in this particularly moist region in the East Himalaya.

## Introduction

Bumble bees (Hymenoptera: Apidae: *Bombus* LATREILLE) are a group of conspicuous, large and colorful bees that mainly inhabit cold and temperate habitats at high latitudes and altitudes. Their conspicuous appearance and abundance made them a prime study object of many early naturalists and insect collectors. After extensive revision in the past decades, around 260 species are currently recognized (Williams 1998; updated online at http://www.nhm.ac.uk/research-curation/research/projects/bombus/index.html).

Current global sampling efforts focus on revising the bumble bee taxonomy at the subgenus level and filling white spots in global distribution data for a worldwide IUCN red list assessment of all species (iucn.org/bumblebees). The latter is urgently needed, since a number of bumble bee species showed dramatic declines in their abundance and range in the recent past (Cameron et al. 2011). The reasons are only partially understood and most likely involve pathogen spillover from commercial breeding, and changes in agricultural practices and land use (Cameron et al. 2011, Jacobson et al. 2018). Moreover, climate change poses a threat to many bumble bee species worldwide, especially those adapted to high altitudes, due to an ongoing decline in suitable habitats (Hoiss et al. 2012, Kerr et al. 2015, Rasmont et al. 2015).

Bumble bees are pollinators of many wild flowers. They are abundant throughout the season and, due to their thermoregulatory abilities, able to be active at very low ambient temperatures (Corbet et al. 1993). Thus, they serve as important pollinators, especially in alpine environments and early in the flowering season (Kevan and Baker 1983, Yu et al. 2012). Besides their ecological importance, bumble bees serve as pollinators for many cultivated fruits, vegetables and spices, making them also economically important. In the industrialized western world, more than one million colonies per year are commercially reared and sold for pollination purposes (Velthuis and van Doorn 2006).

Bumble bees are cold-adapted and therefore are most diverse and abundant in northern temperate habitats and in alpine environments. The Himalaya, the longest mountain range in the world, is home of a high bumble bee diversity due to its variety of suitable habitats. The mountain range spreads over 3,000 km between the Karakorum in the west and the Patkai and Hengduan mountain ranges in the east. As major barrier for the south-eastern monsoon winds, it plays an important role in shaping the climate of entire South Asia (Zhisheng et al. 2001, Xu et al. 2009). The Himalaya is very diverse in local climate, e.g. western end shows strong annual temperature fluctuations and is relatively arid whereas the eastern end is rather stable in the annual temperatures and receives a high amount of annual rainfall. These climatic differences account for distinct differences in flora and fauna (Williams et al. 2010, Rawat 2017). The West Himalaya is characterized by temperate broad leaf forests and arid alpine meadows and pastures at high altitudes with relatively low annual rainfall (Rawat 2017). At the east end, in contrast, annual precipitation can reach up to 5,000 mm (Dhar and Nandargi 2006) allowing the formation of subtropical broadleaf forests and moist alpine meadows at higher altitudes (Rawat 2017). Previous studies found that the biodiversity in the East Himalaya is particularly rich and the region is considered a global biodiversity hotspot (Myers et al. 2000).

So far, bumble bee composition was intensively studied in the West (Williams 1991, Saini et al. 2015) and Central Himalaya (Williams et al. 2010). The highest diversity is reported for the Central Himalaya, from Nepal and the Indian state of Sikkim (Williams 2004, Williams et al. 2010, Saini et al. 2015). Many eastern and western species reach their respective distribution limit in Nepal and the overlap of both faunal regions may contribute to the high bumble bee diversity in this area (Williams et al. 2010). The eastern end of the Himalayan mountain range so far received little attention and only a few actual confirmed records are available (Williams 2004, Saini et al. 2015). The inaccessibility and the climatic conditions make field work in the East Himalaya challenging (see comments in Saini et al. 2015, Rawat 2017) and has certainly contributed to the lack of bumble bee research. Arunachal Pradesh, the northernmost and largest of the Indian north-east region (NER) states, comprises the eastern end of the Himalayan range. Arunachal Pradesh is unique, being densely forested, sparsely populated and agriculturally only extensively managed and thus barely fragmented in its landscape (Tripathi et al. 2016). Previous studies also showed an outstanding biodiversity and high endemism, e.g. in *Rhododendron* species, bamboos, orchids and many other plant taxa (Bhuyan et al. 2003, Mao 2010, Paul et al. 2010, Rawat 2017) and butterflies (Sondhi and Kunte 2016).

In this study, we report the results from the first systematic survey of bumble bees in Arunachal Pradesh, based on material collected during three major and a few minor field trips during the years 2015-2017. The survey represents the first phase of a project aiming at (1) documenting the bumble bee diversity in the East Himalaya to aid global distribution range assessments, (2) identifying local pollinators of fruits, vegetables and crop, and (3) identifying functional adaptations that allow bumble bees to thrive in the particularly challenging climate of the East Himalaya.

## Material & Methods

### Study area and locations

Arunachal Pradesh is the largest of the North-East Indian states and bordered by Bhutan in the west, the People’s Republic of China (Autonomous region of Tibet) in the north, Myanmar in the east and the Indian states of Assam and Nagaland in the south (Fig. 1).

Bumble bee specimens were collected during three major field surveys in the years 2015-2017. The field trips covered the entire flowering season, pre-monsoon (V.-VI. 2016), during monsoon (VIII.-IX. 2017) and post-monsoon (IX.-X. 2015). Additional specimens were collected from the entire state during shorter field visits in the years 2016-2017 (Fig. 1). We covered altitudes between c. 200 m and c. 4,300 m above sea level and habitats ranging from foothill forests (tropical wet evergreen and semi-evergreen), temperate broadleaf forest, subalpine forest up to the alpine zone (Fig. 2, Rawat 2017). GPS locations and altitude were collected using handheld GPS units or cell phones (Garmin Ltd., CH; Apple Inc., CA, USA) and later verified using Google Earth (Google LLC, CA, USA). Mapping of the occurrence data was performed using GPS coordinates and SRTM digital elevation data (Jarvis et al. 2008) in R (R Core Team 2008).

**Figure 1.**
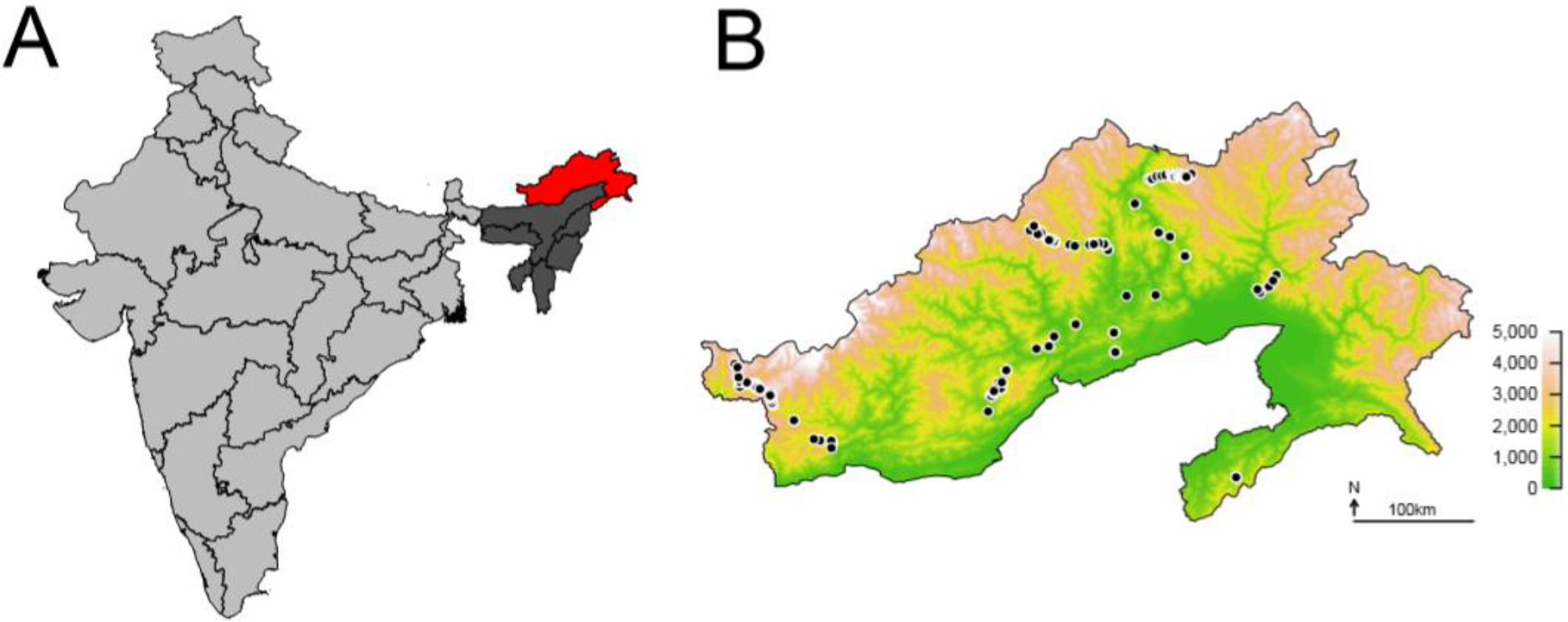
Sampling locations. **(A)** Mainland India with the geographic location of Arunachal Pradesh (red filled area) in the northeast region (NER, dark grey area). Outlines denote Indian state borders. **(B)** Sampling locations within the state of Arunachal Pradesh for three major and a few minor field trips between 2015 and 2017. The locations are projected from GPS data to a SRTM elevation data set. The color scale refers to altitude and does not reflect vegetation zone.

### Sample collection

Bumble bees were collected by sweep netting and immediately killed with cyanide or ethyl acetate. The specimens were then stored in airtight containers with a few layers of tissue and the addition of a few drops of ethyl acetate to prevent mold during the transport. After the field sampling, specimens were dry-mounted on standard insect pins for identification. The collected specimens were deposited in the NCBS Research Collection (National Center for Biological Sciences, Tata Institute of Fundamental Research, Karnataka, Bangalore) for future reference. A full list of the collecting information of the museum specimens is available upon request (curators: Dr. Axel Brockmann and Dr. Krushnamegh Kunte, NCBS Bangalore). In addition to the collected specimens, we also included some field observations. Since these specimens are not available for later reference, we only included specimens that could be unambiguously identified, and from locations where we also collected voucher specimens of the same species. In addition to the specimens collected in this project, we also checked entomological collections for bumble bees from Arunachal Pradesh.

### Experimental ethics

Permits to sample bumble bees were issued by the Government of Arunachal Pradesh to Jharna Chakravorty (No. SFRI/APBB/9/2011-846, No. SFRI/APBB/09/2016/1168) and to Himender Bharti (No. CWL/G/13 (95)/2011-12/Pt./2471-75).

### Species identification

Specimens were identified using published identification keys for adjacent regions, e.g. Kashmir (Williams 1991), Nepal (Williams et al. 2010), Sichuan (Williams et al. 2009), North China (An et al. 2014) and India (Saini et al. 2015). In addition, species’ first descriptions and detailed accounts were consulted (Frison 1933, 1935, Tkalcu 1968a, 1974).

## Results

Between 2015 and 2017, 773 bumble bee specimens were either collected, identified in the field and from photographs or identified in entomological collections (Fig. 1, Tab. 1). 642 specimens were deposited in the NCBS Research Collection. The remaining voucher specimens are part of research project voucher collections (coll. Jaya Narah, Department of Zoology, Rajiv Gandhi University, Itanagar, Arunachal Pradesh) or entomological collections (Department of Entomology, University of Agricultural Sciences, GKVK, Bangalore, India -15 specimens; NBCS Research Collection, Bangalore, India - 2 specimens).

**Table 1.**
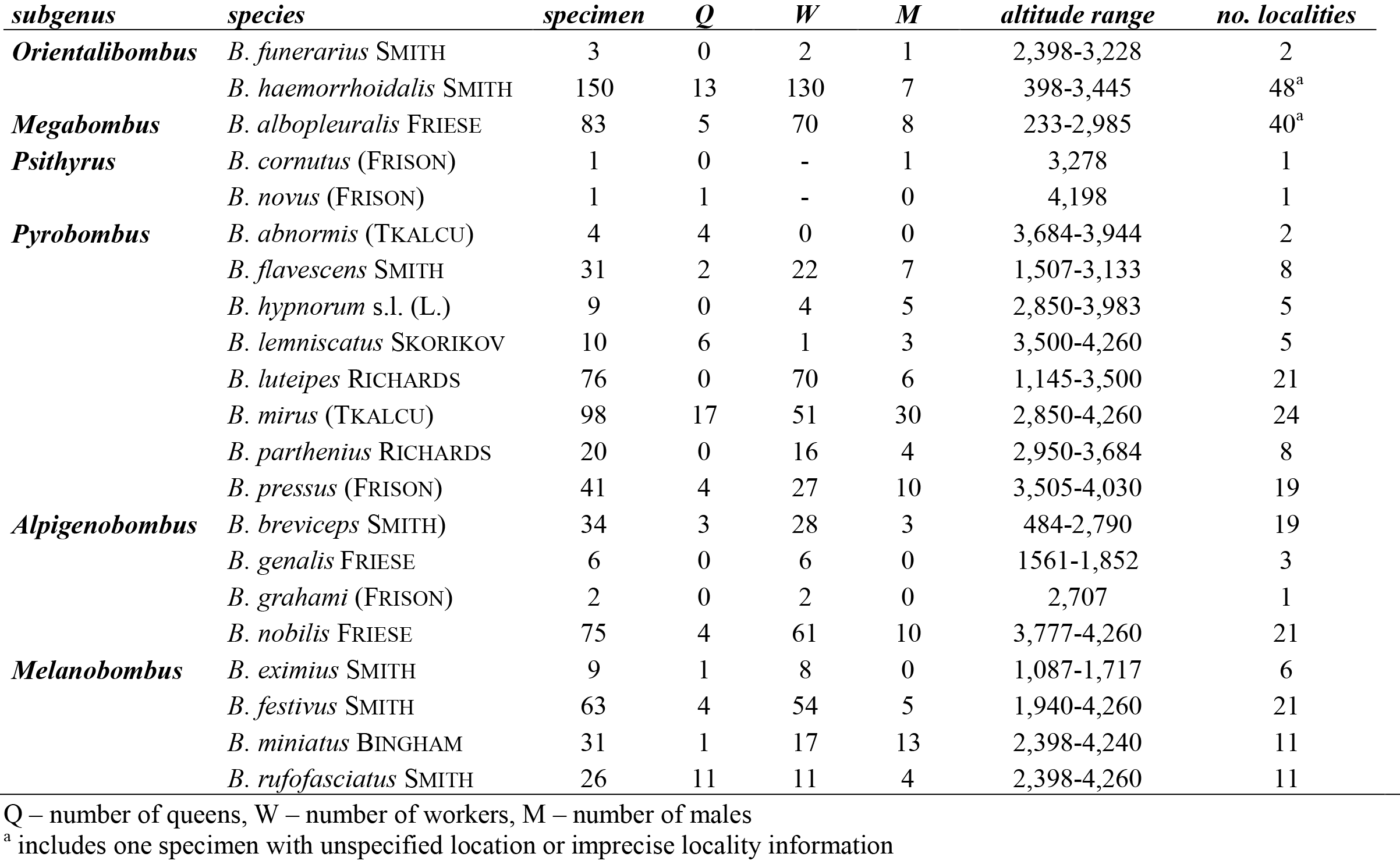
Summary of the collected bumble bee specimens. Listed are all specimens (N = 773) that have been seen and identified by the authors, including material collected during the field trips, specimens from research and museum collections, and specimens identified in the field. Subgenera are sorted according to their phylogenetic position (Williams et al. 2008). Among the subgenera, species are sorted alphabetically

The region sampled covers most of the state Arunachal Pradesh, with less dense sampling in the eastern-most region (Fig. 1). Bumble bees were collected in a large altitudinal range from 233 m to 4,260 m above sea level, covering many different habitat types (Fig. 2). There was a clear altitudinal change in species composition (Fig. 3). In the moist evergreen forest at low altitudes (233-1,085 m), only three species from three different subgenera were observed *(B. haemorrhoidalis* SMITH, *B. albopleuralis* FRIESE, *B. breviceps* SMITH; Tab. 1, Fig. 3, Suppl. Fig. 1B,C,N). Species diversity increased with altitude, climaxing in the region 3,000-4,000 m (mostly corresponding to the subalpine stage) with 15 species from five subgenera (Fig. 3). In total, the specimens belong to 21 currently recognized species from six subgenera (Tab. 1).

**Figure 2.**
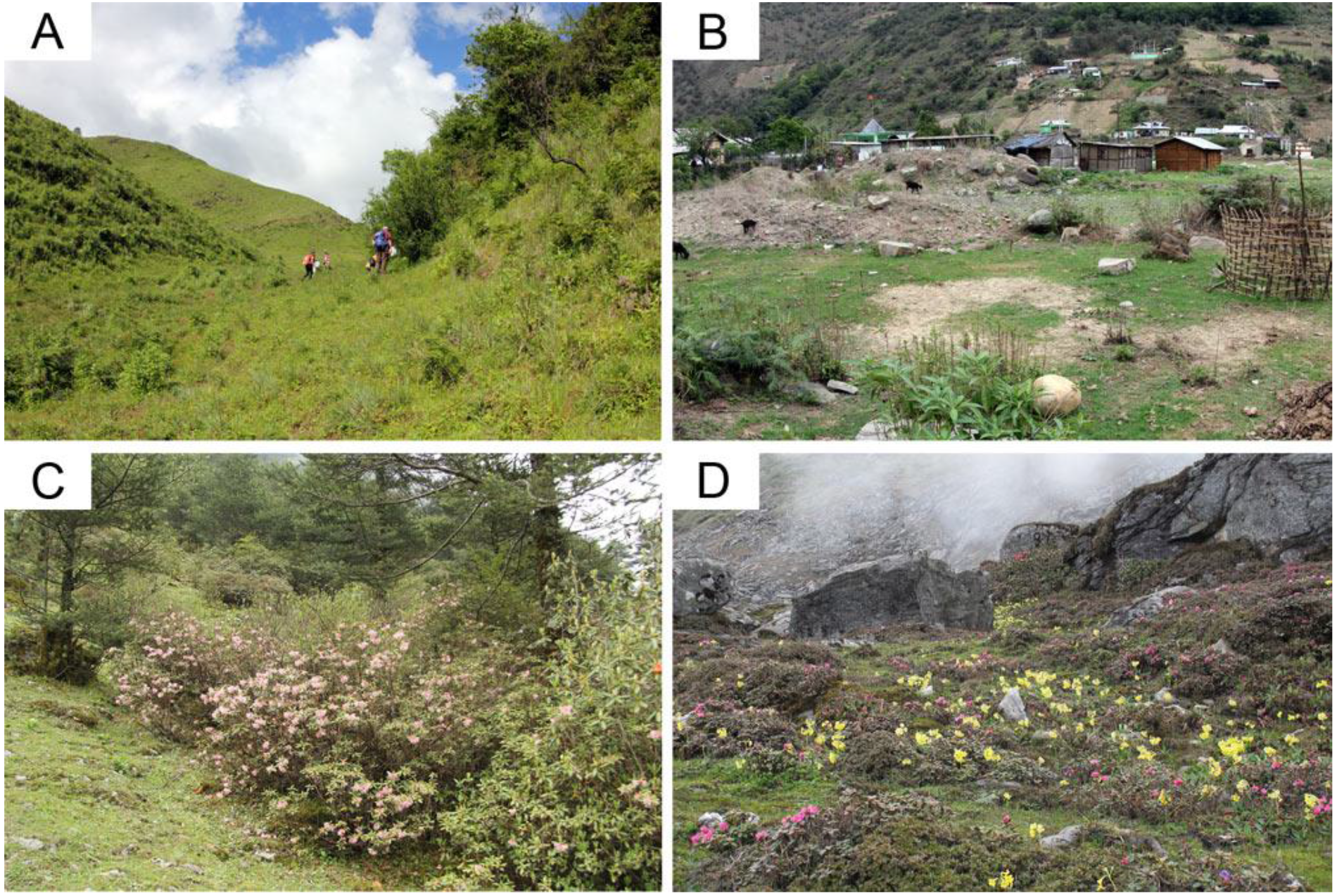
Bumble bee habitats in Arunachal Pradesh. **(A)** Grass-/shrubland at 1,950 - 2,050 m altitude (Mechuka, West Siang district). Workers of *B. festivus* and *B. luteipes* and workers and males of *B. flavescens* were observed visiting *Cotoneaster* bushes. **(B)** Agricultural crops located in a river valley at 1,500 m altitude (Old Dirang, West Kameng district). Workers of *B. flavescens* were collected from *Punica granatum* flowers. **(C)** Ever-green deciduous *Rhododendron-* and *Pinus-* forest at 3,500 m (Karpo, Tawang district), where we collected queens of *B. festivus* and *B. pressus.* **(D)** Alpine meadow with flowering *Primula sp.* and *Rhododendron sp.* (Se-La Pass, Tawang district) at 4,260 m, where we collected *B. mirus, B. lemniscatus*, *B. nobilis*, *B. festivus*, *B. rufofasciatus*, *B. miniatus* and *B. novus.*

**Figure 3.**
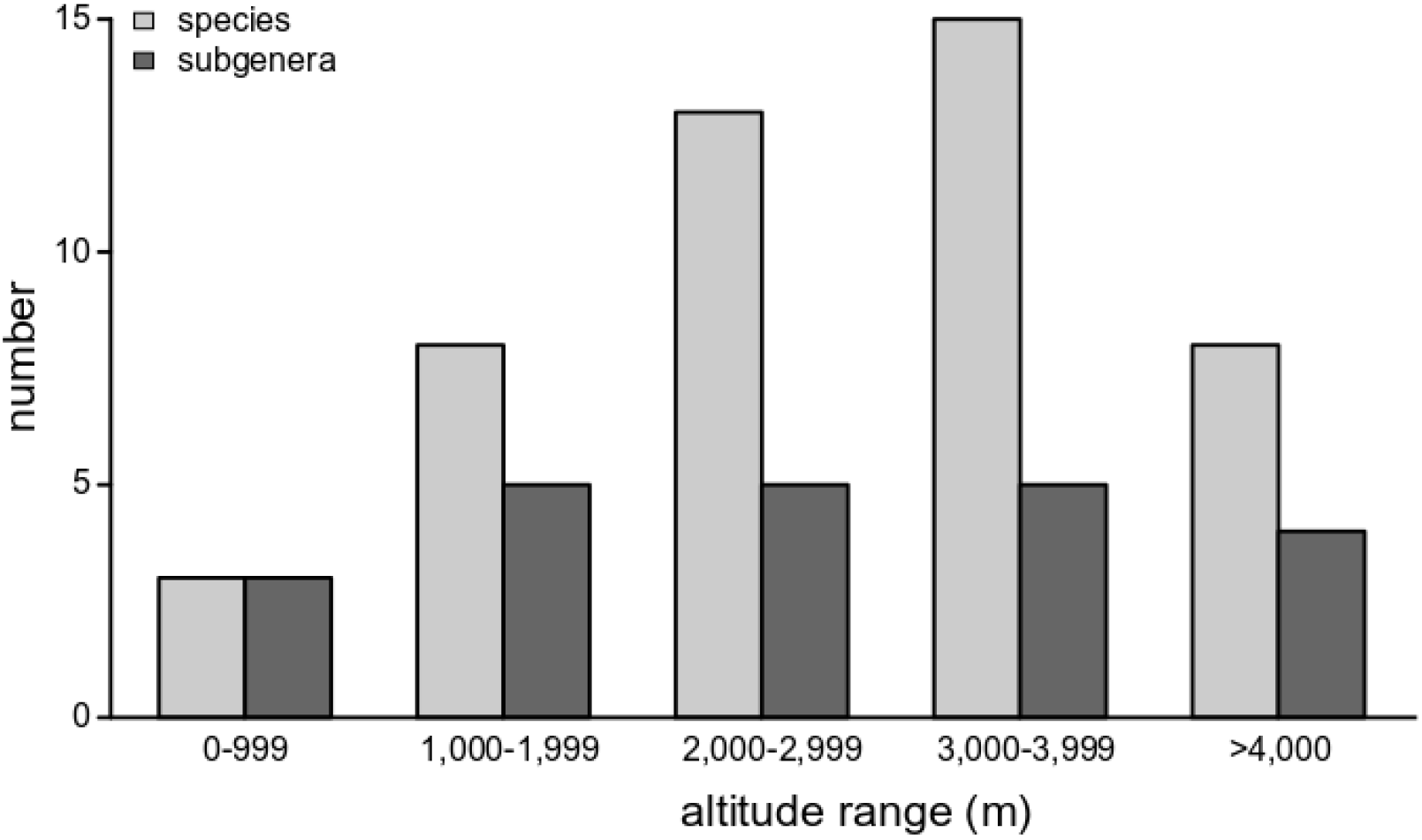
Species and subgenera diversity along the altitudinal gradient. In the lowland tropical forest (<1,000m) only *B. haemorrhoidalis, B. albopleuralis* and *B. breviceps* were observed. With increasing altitude we found an increasing diversity of species. The relatively low diversity at >4,000m may be a sampling bias, since only a few locations were accessible.

## Discussion

### Bumble bee diversity and species records in the East Himalaya

During several field trips in the Indian state of Arunachal Pradesh, we collected > 700 bumble bee specimens belonging to 21 species. This survey represents the first systematic study of bumble bee diversity in the East Himalayan range, which is known as biodiversity hotspot and an important conservation priority region (Myers et al. 2000).

Previously, only very few confirmed records of *Bombus* existed for Arunachal Pradesh. Williams (2004) lists 8 species and predicted the occurrence of another 13 based on their known distribution. During their 12 year survey of India, and based on a total of almost 7,000 specimens, Saini *et al.* (2015) only recorded one species (*B. eximius* SMITH) from this state.

In our study, we collected individuals of 21 currently recognized species (Tab.1), including almost all of the previously confirmed (except for *B. turneri* (RICHARDS)) and more than half of the predicted species (Williams 2004). Furthermore, we collected a number of species that were previously assumed to either have a West (e.g. *B. miniatus* BINGHAM, *B. novus* (FRISON), *B. parthenius* RICHARDS) or Central Himalayan distribution (*B. abnormis* (TKALCU), *B. mirus* (TKALCU), *B. pressus* (FRISON)), and were not expected to occur in Arunachal Pradesh (Williams 2004). Many of these species were previously classified as “vulnerable”, “near threatened” (Williams and Osborne 2009) or “extremely rare” (Saini et al. 2015), are known from a limited number of specimens in entomological collections (P. H. Williams, pers. comment), and could not be found in recent field surveys across India (Saini et al. 2015). *Bombus mirus*, a species previously considered very confined and rare (Tkalcu 1968a, Williams et al. 2010, Saini et al. 2015) even represents ~13% of our entire collection (Tab. 1).

The present list, comprising 22 species (including *B. turneri*, which was not found in our survey), places Arunachal Pradesh close to the species diversity found in the West Himalaya, e.g. Kashmir [29 species], Himachal Pradesh [25] and Uttarakhand [22] (Williams 2004, Williams et al. 2010). Contrary to the East Himalaya, these regions were intensively sampled in the last decades (Williams 1991, Saini et al. 2015). Based on the current sampling status and the predictions by Williams (2004), we expect more species to be found in the future. Alpine regions above the tree line (>4,000m) are scarce and not easily accessible in Arunachal Pradesh (Mishra et al. 2006). A more intense survey of these areas will possibly confirm the presence of high altitude species (e.g. *B. waltoni* COCKERELL, *B. kashmirensis* FRIESE, *B. ladakhensis* RICHARDS, *B. keriensis* MORAWITZ), that are known to occur in South-East Tibet close to the Indian border (Williams 2004, Williams et al. 2015). The East Himalayan region is still vastly under-sampled and more thorough sampling is needed in the entire NER of India at the intersection between the Himalaya and the Patkai mountain range and in the mountain regions of Meghalaya, where the general occurrence of bumble bees is confirmed, but systematic surveys lack (Frison 1933, Tkalcu 1974, 1989, Williams 2004, Saini et al. 2015). Future work in the region will also provide material for taxonomic revisions. Resulting from the large number of specific, subspecific and infrasubspecific synonyms, a genus wide revision is still under progress (Williams 1998). The treatment by Saini *et al.* (2015) not yet incorporates recent taxonomic changes from sub-generic revisions (e.g. Williams et al. 2011, 2012). While the identity of many species in our study is clear from the morphology, a few nominal taxa are currently treated as species complex and future work will likely change their taxonomic treatment (e.g. *B. hypnorum* s.l. (LINNAEUS); see Tkalcu 1974, Williams et al. 2010).

### Mimetic circles

Particularly high local convergence in color patterns is often found within the genus *Bombus.* It is usually interpreted as Müllerian mimicry (Richards 1929, Williams 2007). One of the most remarkable mimetic circles is found in the Himalaya and South-East Asia, comprising *B.(Orientalibombus) haemorrhoidalis*, *B. (Alpigenobombus) breviceps*, *B. (Pyrobombus) rotundiceps* FRIESE and the closely related species of the *B. (Megabombus) trifasciatus*-group (Hines & Williams, 2012; Tkalcu, 1968b; Williams, 1991). The species are members of four different subgenera, corroborating the interpretation that convergent evolution, rather than common ancestry, is responsible for the color pattern similarity.

Three of these species were found in our study area and show identical color pattern across Arunachal Pradesh. Two other mimetic groups are present in the region, each comprising members of at least two different subgenera. First, *B. (Pyrobombus) abnormis*, *B. (Pyrobombus) hypnorum* s.l. and workers of *B. (Melanobombus) festivus* SMITH all have a brown thorax and a white tail. The second circle comprises *B. (Pyrobombus) flavescens* SMITH, *B. (Melanobombus) eximius* and *B. (Alpigenobombus) genalis*, which are characterized by black body pile, orange tinted wings and orange-brown cuticle and hairs on the legs (see examples in Fig. 4). Color pattern convergence within *Bombus* is also often observed between the parasitic species of the subgenus *Psithyrus* and their preferred host species (Reinig 1935, Williams 2008). The parasitic *B. novus* (FRISON), recorded in our study, was previously assumed to develop in nests of *B. rufofasciatus* SMITH (Tkalcu 1974). Although the female of *B. novus* shares with *B. rufofasciatus* a reddish band of pile just anterior to the white tail, it more closely resembles *B. miniatus* in the pale yellow (rather than white-grey) coloration of the anterior pale bands and the darker tint of the wings (Williams et al. 2010; Suppl. Fig 2). Furthermore, the known distribution ranges of the latter match more closely, being (mostly) West Himalayan species that both reach their eastern distribution limit in Arunachal Pradesh, whereas *B. rufofasciatus* is a widespread Himalayan and Tibetan species (Williams et al., 2015). However, most *Psithyrus* are to some extent flexible in their host choice and more observations, especially from breeding *Psithyrus* in their host nests, are necessary to confirm this suggested parasite-host association (Williams, 2008).

**Figure 4.**
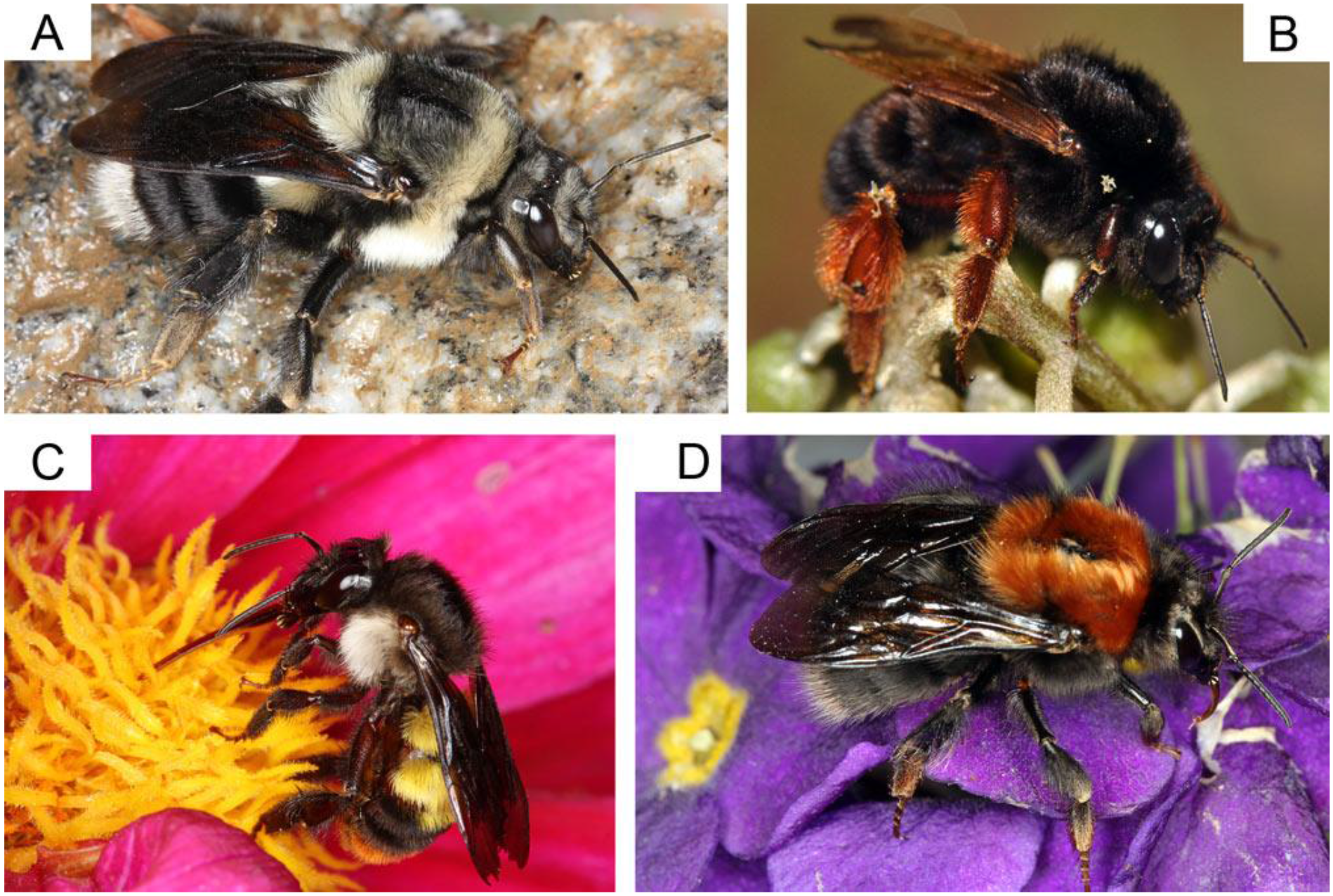
Examples of bumble bee specimens collected in Arunachal Pradesh. **(A)** *Bombus miniatus* (queen) is a West Himalayan species of the subgenus *Melanobombus* reaching its eastern distribution limit in Arunachal Pradesh. **(B)** *Bombus genalis* (worker), a rare species of the mid-elevation that is narrowly distributed in the East Himalaya. **(C)** *Bombus albopleuralis* (worker), a wide-spread Himalayan species that occurs in a large altitudinal range from the tropical lowlands to the subalpine zone in Arunachal Pradesh. **(D)** *Bombus abnormis* (queen), an elusive and very rare high elevation species of the subgenus *Pyrobombus* that is narrowly distributed in the East Himalaya.

### Altitudinal distribution and adaptation

Covering a large range of altitudes and habitat types, we found clear patterns of species-specific altitude ranges (Fig. 3). A number of species are only found in the subalpine and alpine region at the highest elevations, and they occupy similar altitude niches as in other regions of the world (e.g. *B. abnormis, B. lemniscatus* SKORIKOV, *B. mirus, B. nobilis* FRIESE, *B. pressus* (FRISON); Williams et al. 2009, 2010). We observed the highest species diversity at altitudes between 3,000-4,000 m (Fig. 3), similar to observation in the Central Himalaya (Williams et al. 2010). However, at the current stage, this may also represent a sampling bias from the relatively lower number of sampling points at high altitudes. In general, species diversity was found to decline towards lower elevations and in the lowland (<1,000 m) only three species (*B. haemorrhoidalis, B. albopleuralis, B. breviceps*) were found. These species also occur at relatively low elevations throughout the Himalaya (lowest elevations: *B. haemorrhoidalis:* Kashmir - 1,000 m, Nepal - 850 m, *B. albopleuralis:* Kashmir - 1,000 m, Nepal - 950 m, *B. breviceps:* Nepal - 980 m; Williams 1991, Williams et al. 2010). Our records (*B. haemorrhoidalis* - 398 m, *B. albopleuralis* - 233 m, *B. breviceps* - 484 m; see Tab. 1), represent the lowest elevations at which these species, and bumble bees in general, have ever been recorded in the Himalayan range (Williams 1991, Williams et al. 2010). Bumble bees often occur in a wide altitudinal range, but only few species reach the tropical lowland, where conditions are usually unfavorable for these cold-adapted bees (Moure and Sakagami 1962, Williams 1991, Gonzalez et al. 2004, Williams et al. 2009).

Our observation may have multiple, not mutually exclusive, explanations. First, the specific climate of the East Himalaya probably allows certain bumble bee species to thrive at relatively lower altitudes (see below). Indeed, there seems to be a gradual decrease in the lower elevation limit from the west to the east that supports this interpretation (Williams 1991, Williams et al. 2010). Second, bumble bee workers can cover large horizontal and, particularly in steep terrain, vertical distances during their foraging trips (Osborne et al. 1999). In Arunachal Pradesh, most of the valleys are particularly steep and both lowland and higher elevations are within the foraging distance of a few kilometers. Therefore, the low records may represent foraging workers from a nest at higher altitude.

*B. haemorrhoidalis, B. albopleuralis* and *B. breviceps* cover a wide range of altitudes and usually were most abundant at medium elevations (Tab. 1, Suppl. Fig. 1). Nevertheless, the wide range of foraging habitats, each posing their own challenges with respect to thermoregulation and energy expenditure, is remarkable. Future work is necessary to assess their specific individual and population-level adaptations that provide the plasticity to cover such a diversity in altitudes and habitat types, while other species are restricted to very small ranges and specific habitats (Williams et al. 2009, 2010, 2018). This plasticity (or absence of) is of particular interest when we want to understand potential threats due to climate change, making some species more vulnerable than others.

Several physiological and behavioral adaptations have been discussed in the context of altitudinal adaptation in bumble bees and previous work shows that behavioral plasticity allows quick adaptation to different altitudes (Dillon et al. 2006, Dillon and Dudley 2014). At the morphological and physiological level, wing load and wing aspect ratio (Cartar 1992), variation of the cuticular hydrocarbon composition, which prevents bees from desiccation (Foley and Telonis-Scott 2010, Menzel et al. 2017), or changes in mitochondrial density and/or enzyme composition (Harrison et al. 2006, Zhang et al. 2013) may be important factors that vary among populations. However, the specific adaptations that allow these species to thrive in the particularly challenging habitats in the East Himalaya, where the peak of the monsoon season coincides with the peak of colony development in many species, is subject to future investigations. Our survey identified *B. haemorrhoidalis* and *B. albopleuralis* as suitable model taxa to investigate the potential adaptations to specific climatic conditions at the individual and population level. Both species cover a wide range of altitudes and are widely distributed in Arunachal Pradesh (Tab. 1. Suppl. Fig. 1).

### Current and Future Threats and Conservation

The finding of many rare and confined species of bumble bees in Arunachal Pradesh highlights the importance of extensive sampling in remote regions to better understand species distribution and ecological requirements (see also the discussion in Williams 2018).

Although many species may be confined or rare from a global perspective, they can be locally abundant and/or restricted to a very specific habitat. The specific climate of the East Himalaya, with the high amount of precipitation, supports a high biodiversity including a large amount of endemism in the region (Myers et al. 2000, Mao 2010). Our observations suggest that a couple of bumble bee species may be particularly adapted to these conditions as they are restricted to a very limited region in the East Himalaya (e.g. *B. mirus, B. genalis*).

Arunachal Pradesh can currently be considered a remote region without serious recent land use changes, only small scale agriculture and a very low population density (Sikri 2006). However, locally distributed species and high altitude specialists may still be under future threat of extinction, through changes in agricultural practices or climate change (Xu et al. 2009, Hoiss et al. 2012). Rising temperatures force bumble bee species to shift to higher elevations (Kerr et al. 2015), but high elevation refuges may be limited for species that are adapted to the East Himalayan climate. It is therefore crucial to understand the adaptations of the local bumble bee fauna to assess their future threat status. Furthermore it is crucial to develop general strategies for the future to preserve much of this remarkable region (Myers et al. 2000, Anonymous 2011).

In the Himalaya, bumble bees serve as important pollinators of many fruits and vegetables, e.g. cardamom (Deka et al. 2011), apple and other fruit (Raj et al. 2012, Raj and Mattu 2014) and crop (Tayeng and Gogoi 2018). Understanding their ecological requirements and preserving the habitats to support pollinator diversity is crucial for a sufficient agricultural yield, especially in the extensively managed small-holder farming systems that are abundant in Arunachal Pradesh (Kala 2005). Bumble bees are used worldwide as pollinators for commercial fruit and vegetable production (Velthuis and van Doorn 2006). Initially, commercially reared species were used outside their native range, resulting both in the introduction of alien species (Morales et al. 2013) and pathogen spread to native bumble bee populations (Arbetman et al. 2013). Nowadays, attempts are made to select suitable native species and develop methods for their commercial rearing in many world regions (Padilla et al. 2017). Laboratory rearing of *B. haemorrhoidalis* in India (Chauhan et al. 2014) and *B. breviceps* in Vietnam (Thai and Van Toan 2018) are first steps to produce native bumble bee colonies for commercial pollination. Both species are widespread in Arunachal Pradesh and would make good pollinators for many fruit and vegetables (Deka et al. 2011). Additional work is now necessary to either confirm its potential or find other promising species for the future development of commercial fruit and crop pollination in Arunachal Pradesh.

## Acknowledgements

We are particularly grateful to the students that helped during the field collections (Rajiv Gandhi University: Mosses Messar, Mohin Raza Naqvi, Nyaton Kitnya; Punjabi University: Joginder Singh, Sishal Sasi; University of Würzburg: Franziska Bandorf) to Tapir Darang, and the drivers and carriers who made the field trips possible.

We further thank the curators of the NCBS Research Collection (Dr. K. Kunte) and the Collection of the Entomology Department of the University of Agricultural Sciences, GKVK, Bangalore (Prof. V. Belavadi) for access and support during our visits to the collections and the permit to use their data for our distribution records. P.H. Williams (National History Museum, London, UK) provided much valuable information and kindly identified a few voucher specimens.

The bumble bee mapping project is part of the Chemical Ecology Network Programme funded by the Department of Biotechnology, Govt of India (Sanction number-No. DBT-NER/Agri/24/2013). The Field trips of researchers from RGU and NCBS were supported by funds from Chemical Ecology Project. Researchers from the University of Würzburg and Vienna were supported by institutional funds from the University of Würzburg to Thomas Schmitt and Johannes Spaethe.

